# Cellulose Rich Food Leads Anxiety through Gut-Brain Axis-mediated Amygdalar Dopamine Upregulation

**DOI:** 10.1101/2024.05.07.593082

**Authors:** Kaede Ito, Haruka Hosoki, Yuya Kasai, Hiroyuki Sasaki, Atsushi Haraguchi, Shigenobu Shibata, Chihiro Nozaki

## Abstract

It is widely said that healthy intestinal environment takes essential role for better mental condition. One of the known dietary nutrients which maintains intestinal environment is the dietary fiber. Recent study showed that maintaining intestinal environment by dietary fiber succeeded to alleviate the psychiatric disorder symptoms in animals. However, such effects have only been reported with soluble fiber, which is highly fermentable and promotes short-chain fatty acid (SCFA) production, and not with insoluble fiber. Therefore, we aimed to verify whether insoluble fiber, such as cellulose, can alter emotion via changes in the gut. We divided mice into two groups and fed either standard diet (SD, contains both insoluble and soluble dietary fibers) or cellulose rich diet (CRD, contains cellulose alone as the dietary fibers). The CRD-fed mice displayed 1) the increased the anxiety-like behavior accompanied with 2) the modified amygdalar dopamine signaling. We further found the decreased intestinal SCFA levels along with intestinal permeability, dysmotility and hypersensitivity in CRD-fed mice. These behavioral and physiological effect of CRD has been completely abolished in vagotomized mice, indicating the direct link between intestinal environment exacerbation to the emotion through gut-brain axis. Additionally, the opioid antagonist abolished the CRD-induced anxiety, suggesting the involvement of opioidergic system to the anxiety which may evoked by increased amygdalar dopamine levels. Altogether, our findings suggest that consumption of cellulose alone as the dietary fiber may evoke intestinal abnormalities which fires the vagus nerve then opiodergic system and amygdalar dopamine upregulation, resulting in the enhancement of anxiety.

**Graphical Abstract: Possible mechanism of CRD-induced anxiety unveiled by current study:** Our study clarified that long-consumption of cellulose-rich food (CRD) will lead decrease of SCFAs which may cause the intestinal disability, including decreased motility and increased intestinal permeability as well as upregulation of TRPA1 and SGLT1. These physiological modifications resulted as the intestinal hypersensitivity, which possibly overstimulate the vagal transmission which may activate endogenous opioidergic systems such as enkephalin (Enk) at the nucleus tractus solitarii (NTS). The activation of opioidergic system may suppress the GABAergic neuron in ventral tegmental area (VTA), resulting in the excess release of dopamine and further receptor modification in amygdala (Amyg), which might in the end cause the characteristic anxiety. The figure was created with BioRender.com.

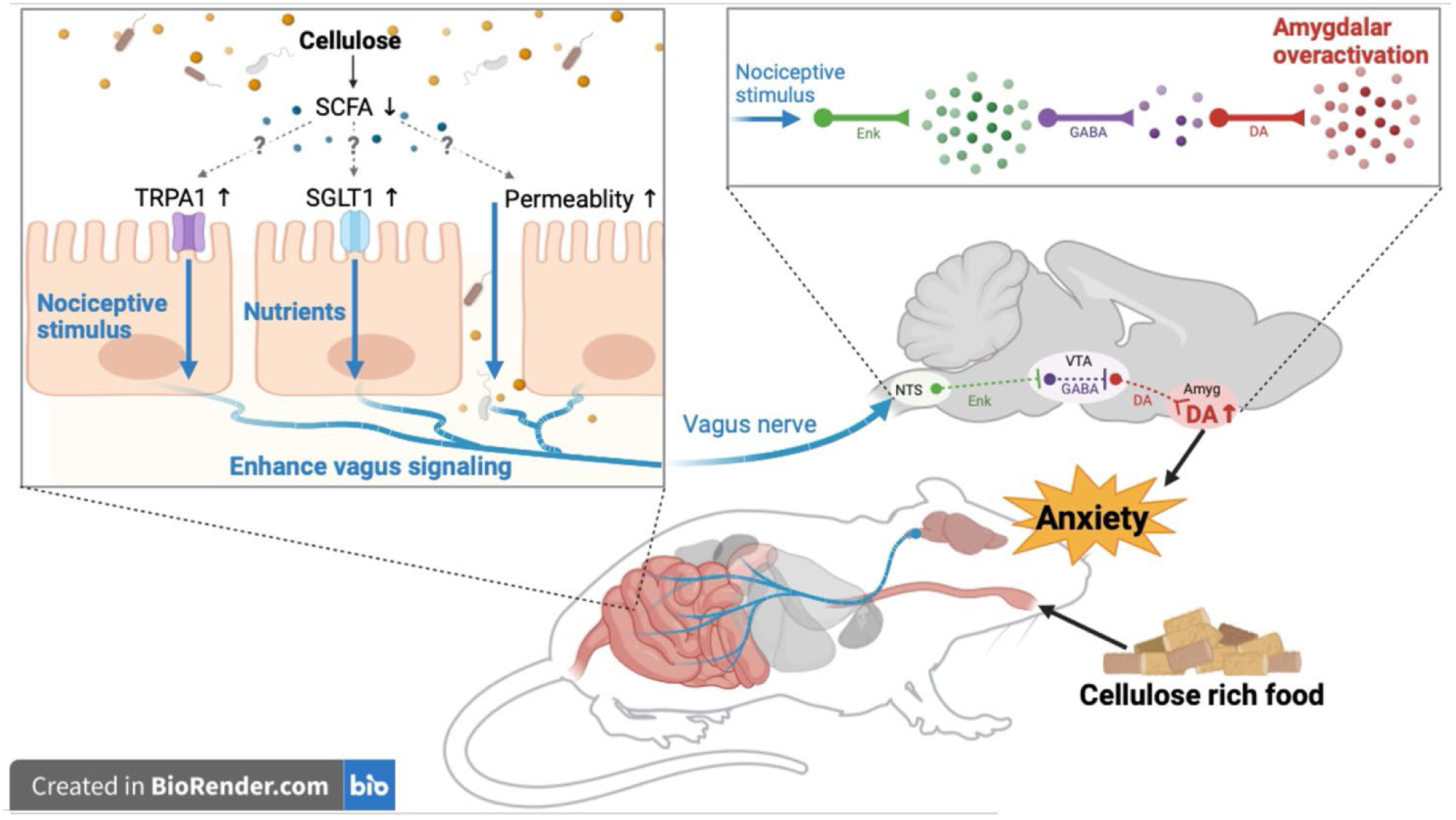

## Introduction

In recent years, the number of people suffering from daily stress and anxiety as well as mental disease has been increasing. WHO has reported that in 2019 there were 970 million people suffering mental disease worldwide, which is more than 10 % global population(1). Among the numerous diseases, mental disorders were the second most prevalent disease of Years Lived with Disability worldwide in 2019(2), showing that mental illness is a long-term affliction for many people. However, the diagnosis and treatment of mental illnesses are fraught with difficulties. The diagnosis for psychiatric disorders has unresolved issue like distinguishing risk from disorder, which makes difficult to select appropriate treatment (3,4). Further, the response rate of both psychotherapies and pharmacotherapies for psychiatric disorders is ≤50%, suggesting insufficient development of effective treatments(5). These issues call for elucidation of the pathophysiology and pathogenesis of psychiatric disorders, and further development of therapeutic methods(6).

Recently, the relationship between the intestinal environment and psychiatric disorders has been attracting attention(7). Intestinal environment of depressed patients is known to deteriorate(8), whereas germ-free mice display the resistance to the depression(9). These reports strongly suggests that the maintenance of healthy intestinal environment and microbiota may improve mood and emotion. Even the clinical trials in depressed patients have shown that probiotic intake alleviates anxiety along with the gut microbiota modification(10). It has also been shown that administration of probiotics alters the GABAergic nervous system, resulting in an anxiolytic effect in Native mice(11).

Although probiotics are the most supported “intestinal health regulators”, dietary fiber also play the important role of maintaining healthy intestine(12). Dietary fiber is indigestible and is digested by intestinal bacteria, promoting the production of short-chain fatty acids (SCFAs). SCFAs are known to play an important role in maintenance of intestinal homeostasis, such as suppression of inflammation and intestinal immunity, suppressing various diseases(13). However, these beneficial effects vary widely among different types of dietary fiber. Dietary fiber is roughly classified into two categories based on its solubility in water: insoluble fiber and soluble fiber. We previously showed that the SCFA concentrations in the cecum contents was increased in soluble fiber (digestion resistant dextrin)-fed mice, compared with those of insoluble fiber (cellulose)-fed animals(14). Soluble fiber is fermentable and has a stronger intestine-regulating function, therefore is used as prebiotics(15). Number of past studies suggest that soluble dietary fiber alleviate psychiatric disease through improvement of the intestinal environment. Thus, improvement of the intestinal environment by soluble fiber treatment alleviates symptoms of depression(16), schizophrenia(17), and diabetes-induced anxiety(18) in rodents. However, on the other hand, the effects of insoluble fiber such as cellulose, on the intestinal environment and emotions have not been examined at all.

Cellulose is the main constituent of plants and therefore the most common dietary fiber consumed daily. Since cellulose is less fermentable than soluble fiber yet has high water retention properties, it can increase the bulk of stool and physically stimulate the intestines to improve peristalsis (bowel movements)(19), even in chronic constipation patients(20). High-cellulose diet also was reported to have protective effect against dextran sodium sulfate-induced colitis by modulating lipid metabolism and gut microbiota, suggesting that cellulose is effective in maintaining intestinal homeostasis(21).

We therefore aimed to verify whether insoluble fiber can alter emotion via changes in the gut. Strikingly, we found that long-term cellulose rich food (CRD) modified the dopamine signaling in amygdala which led the rise of anxiety-like behavior. Such emotional and neuronal modification might be due to intestinal hypomotility, and hypersensitivity caused by SCFA decrease by CRD exposure, which was ameliorated by either vagotomy or inhibition of opioid receptors. Thus, our results suggest that CRD-induced intestinal changes evoked amygdalar dopamine signaling abnormalities via the vagus nerve then opioidergic system, which in turn lead the anxiogenic effect.

## Results

### The long-term ingestion of cellulose rich food enhanced the anxiety-like behavior in mice

Mice were divided into two groups and fed either standard diet (SD: contains 2.8% both insoluble and soluble dietary fibers) or cellulose rich food (CRD, contains 5% cellulose). Then we assessed if CRD effects on the anxiety level in mice by behavior tests (Figure 1 and Figure S1). We found that CRD-fed mice display the increase of anxiety-like behavior in marble burying test (Figure 1C, D) without the change of locomotor activity (Figure S1). On the other hand, there were no significant differences in anxiety-like behavior in open field test and elevated plus maze test (Figure S1). The level of corticosterone, a typical stress hormone, tended to be elevated, but was low in the urine of 16-week CRD-fed animals (Figure 1E, F). These results indicates that the long-term ingestion of CRD time-dependently enhanced the anxiety level and stress response in mice.

**Figure 1:**
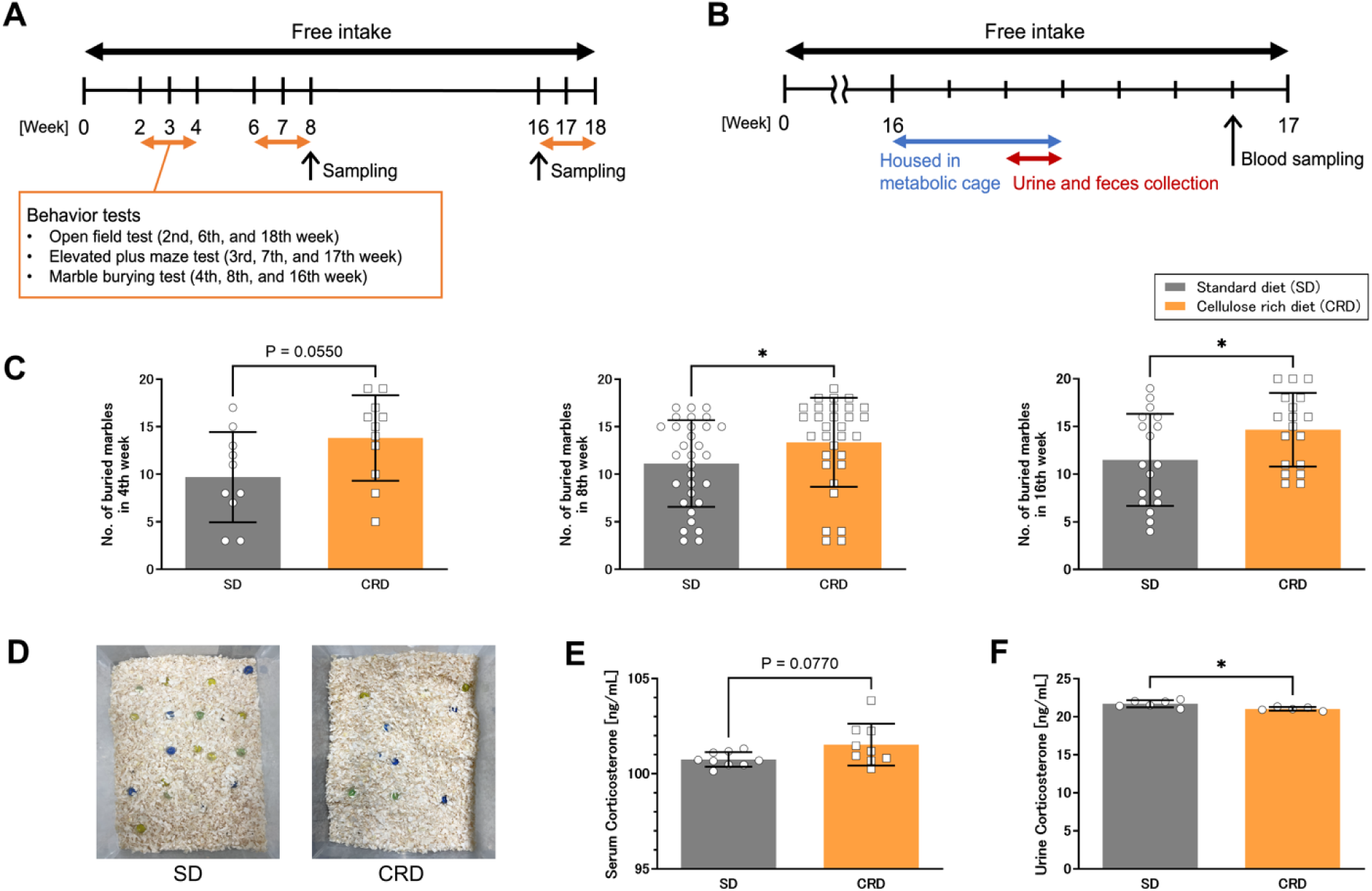
CRD-fed mice displayed the increased anxiety level and the corticosterone level modification. (A, B) Overall scheme of the experiments. Mice were divided to two group and were fed either standard diet (SD) or cellulose rich diet (CRD). Behavioral tests have been done at 2-4th, 6-8th and 16-18th week of feeding period with one-week interval for each testing. Individual group of animals are prepared and used for other testing and sample collection at 8th and 16th week respectively. Feces and urine collection Serum and urine for corticosterone measurements were sampled from mice exposed to CRD for 16 weeks. Mice were housed in the metabolic cage for three-days then feces and urine samples are collected 24 hours at third day (B). (C) CRD-exposure transiently increased the number of buried marbles on the marble burying test. (D) The example of marble burying observed at 16th week. CRD animals buries more marbles than SD groups. (E, F) The corticosterone concentrations in serum and urine are measured by ELISA. While CRD tend to increase serum corticosterone, it significantly decreased the urine corticosterone. Data are expressed as means±SEM of 10-31 mice/group for behavioral tests, and 5-9 mice/group for corticosterone measurement. Statistical significant is indicated by \**p* < 0.05 according to unpaired t-test or Mann Whitney test.

### The long-term ingestion of cellulose rich food modified the dopaminergic system and proinflammatory cytokine release in amygdala

To understand which molecular mechanisms underlying CRD-induced anxiety, we measured the monoamine level in various brain regions that is known to involve in the regulation and expression of anxiety. We found that 16-week CRD-exposure significantly increased the dopamine level in amygdala (Figure 2B), while no significant differences were seen in other brain regions or for other monoamines (Figure 2A, Figure S2A, B). Interestingly, dopamine level of striatum and nucleus accumbens had no change despite of dense dopaminergic neuron projection(22) (Figure S2C, D), indicating that CRD may evoked modification of dopaminergic system specifically at amygdala. We therefore measured the mRNA expression levels of dopamine receptor and transporter in amygdala. Interestingly, CRD-fed mice displayed the significant lower mRNA expression of dopamine D2 receptors (D2R) compared the SD-fed mice in the 8th week of ingestion period (Figure 2B), whereas there were no significant differences in 16th week (Figure 2C).

**Figure 2:**
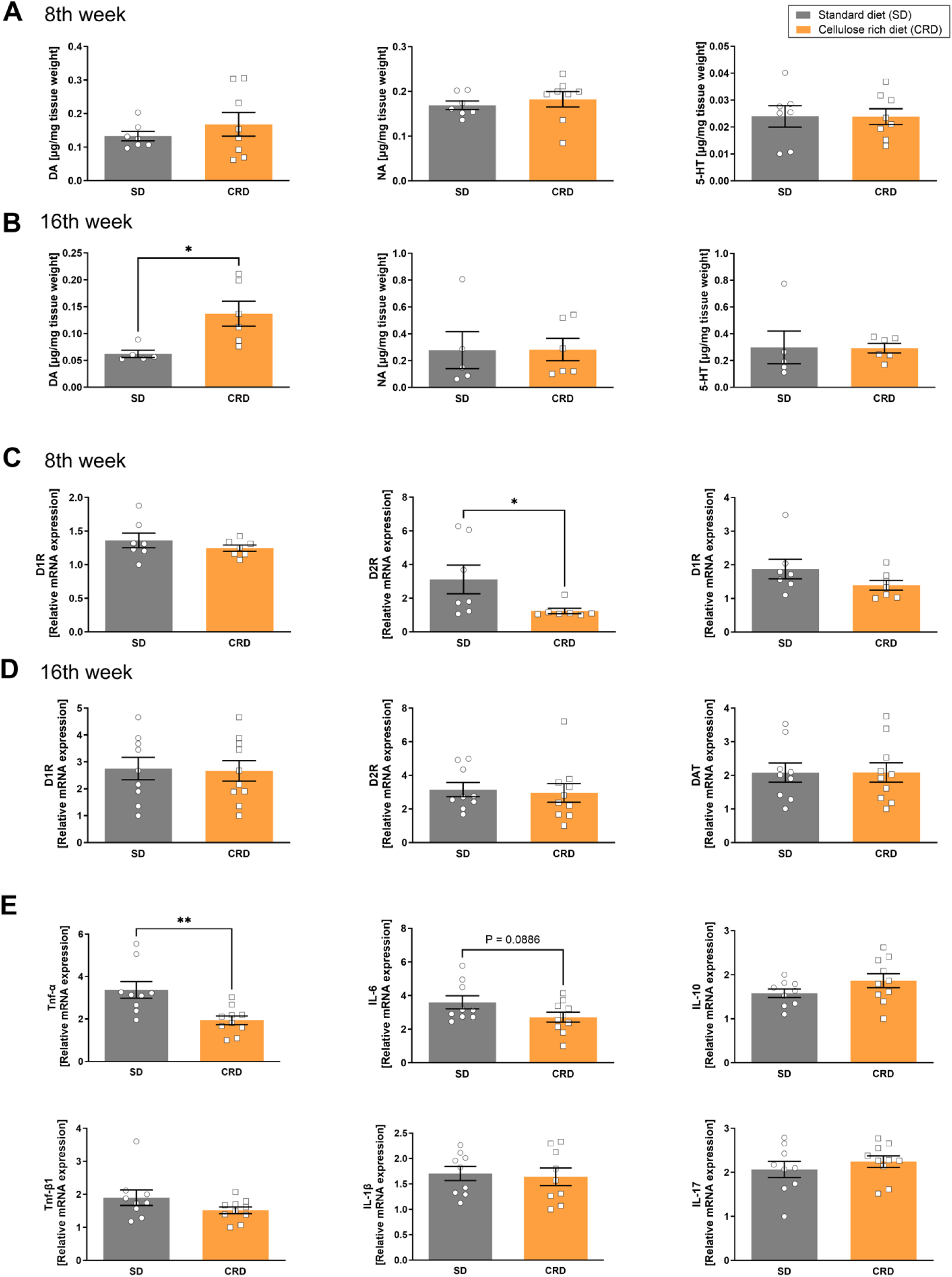
Chronic CRD modified dopamine signaling and proinflammatory cytokine release in amygdala. (A, B) Dopamine (DA), noradrenaline (NE), and serotonin (5-HT) level in amygdala was measured by HPLC and compared between SD- and CRD-fed mice at feeding period of 8 weeks (A) and 16 weeks (B). (C, D) The amygdalar mRNA expression level of dopamine receptors (D1R and D2R) as well as dopamine transporter (DAT) was determined at 8th (C) and 16th (D) week after the exposure to either SD or CRD. (E) The amygdalar mRNA expression level of cytokines at 16th week after the exposure to either SD or CRD. Data are expressed as means±SEM of 5-10 mice/group. K

As recent studies suggests that neuroinflammation might contribute to the emotional change(23), we also focused on the amygdalar inflammation. To evaluate the inflammation, we measured the mRNA expression levels of pro- and anti-inflammatory cytokines in amygdala. Surprisingly, the mRNA expression levels of amygdalar TNF-α and IL-6 was decreased in CRD-fed group (Figure 2D). Flow cytometric analysis of bone marrow also showed the decrease on monocyte population (Figure S3), which may suggest that CRD exposure decreased monocyte production in bone marrow to decrease the inflammatory cytokine levels in amygdala.

### Vagal signaling mediates CRD-induced anxiety

Next question will be whether CRD directly or indirectly modulated amygdalar signaling to have anxiogenic effect. Since the gut-to-brain transmission mediated by the vagus nerve has been suggested to modulate the anxiety(24,25), we conducted vagotomy, the transection of hepatic vagal branch (Figure 3A). If CRD-evoked anxiety will be diminished by vagotomy, it demonstrates that CRD do not directly modulate amygdala, but rather indirectly modify it possibly through gut-brain axis. Consequently, vagotomized animals did not display any sign of CRD-induced anxiety (Figure 3B) with no change of motor ability (Figure 3C). Vagotomy also tended to reverse CRD-induced decrease in D2R expression (Figure 3E), suggesting that vagal signaling is involved in changes in the amygdalar dopaminergic system. However, vagotomy had no significant effect on CRD-induced increases in amygdalar dopamine level (Figure 3D).

**Figure 3:**
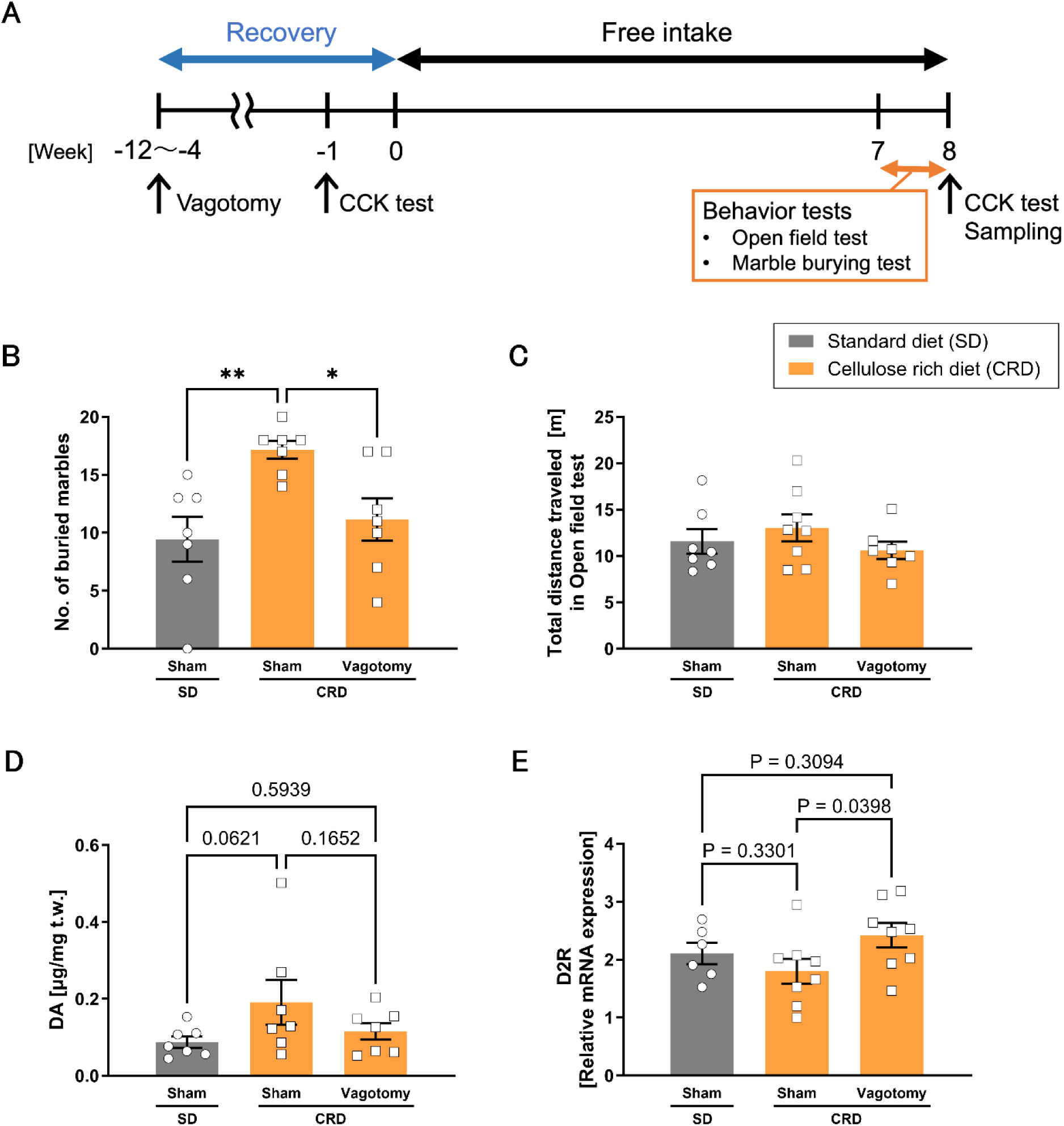
CRD-induced anxiety and dopaminergic modulation has been suppressed by vagotomy. (A) Overall experimental scheme for the vagotomy. At least 4-weeks of recovery time was taken after the surgery. Then mice were fed with standard diet (SD) or cellulose rich food (CRD) for another 8-weeks, then behavioral tests were performed, and sacrificed to collect the tissue. (B, C) CRD-evoked anxiety-like behavior in marble burying test was suppressed by vagotomy (B) with no effect on locomotive activity (C). (D) Vagonomy had no significant effect to the amygdalar dopamine level. (E) Decrease of amygdalar D2R mRNA expression level in CRD animals has been reversed by vagotomy. Data are expressed as means±SEM of 7-8 mice/group. Statistical significant is indicated by \**p* < 0.05, \*\**p* < 0.01 according to one-way ANOVA with uncorrected Fisher’s LSD.

These results show that gut-brain axis composed by vagus nerve is essential for the CRD-induced modification of amygdalar dopamine signaling. It also indicates that anxiogenic effect of CRD has happened through the modification of intestinal environment rather than direct modulation of brain.

### CRD caused worsening of intestinal environment including increased intestinal permeability and dysmotility

As suggested above, CRD-induced anxiety is mediated through vagus nerve possibly from intestinal input. We hence evaluated what intestinal modification CRD may cause. The first sign of intestinal modification we found in CRD-fed animals was the significant increase of cecum pH (Figure 4B). This could be explained by the further measurement of intestinal short chain fatty acid (SCFA) and lactic acid contents, which has been remarkably decreased by CRD consumption (Figure 4C and D). Importantly, vagotomized CRD-mice that displayed improvement of anxiety and amygdalar dopamine signaling (see Figure 3) also showed significant increase of cecum pH together which may be led by decrease of SCFAs (Figure S4). These results suggest that long-term CRD consumption may cause the decrease of SCFA production in intestine, which may lead to higher intestinal pH, one of the hallmark of worsened intestinal environment(13).

**Figure 4:**
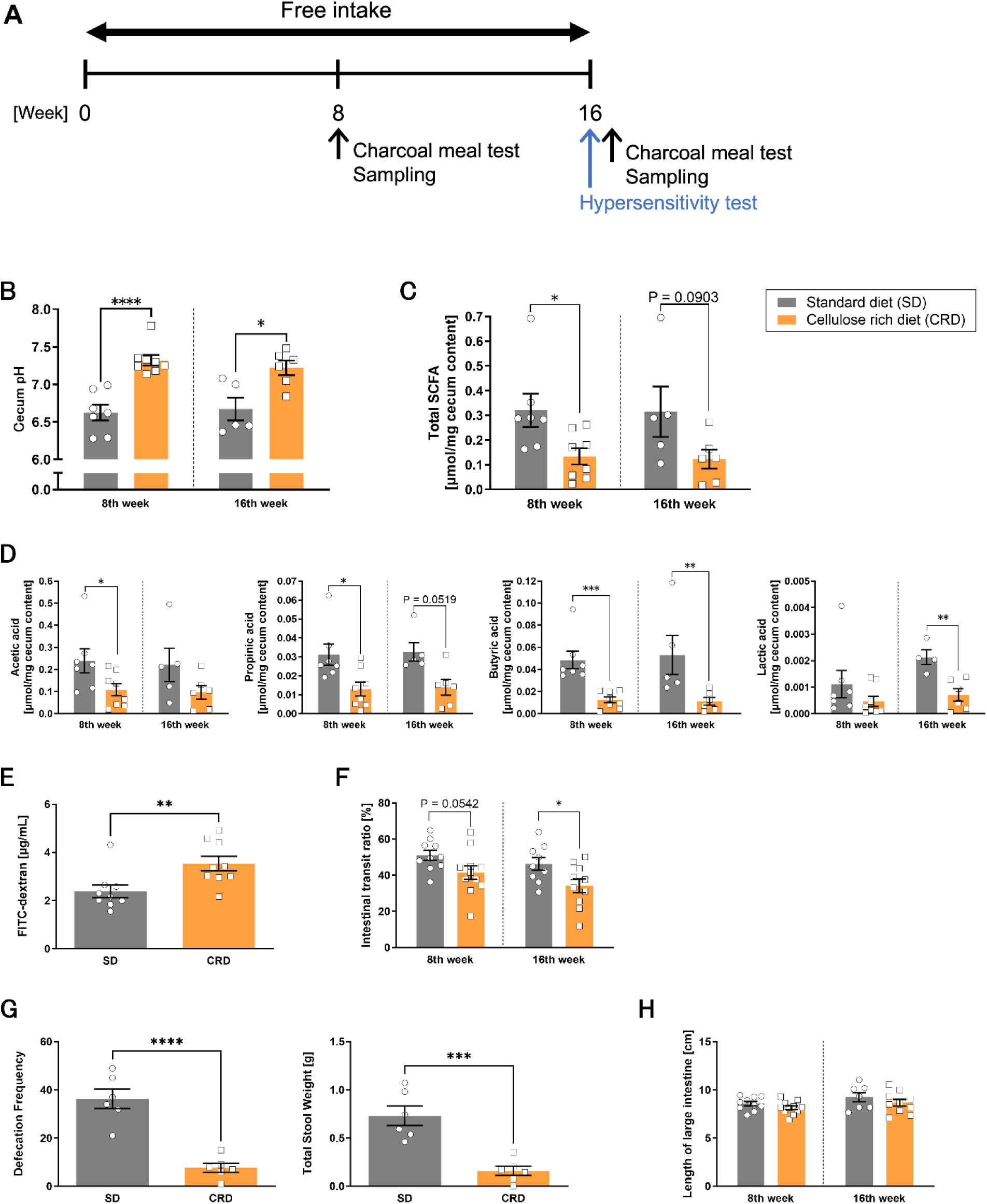
CRD consumption evoked the remarkable deterioration of intestinal environment. (A) Overall scheme of the experiments. Individual group of animals are prepared and used for other tests and sample collection at 8th and 16th week respectively. (B-D) Increase of cecum pH (B) which may be caused by significant decrease of SCFA and lactic acid in cecum content (C, D) has been observed in animals which consumed CRD for 8-weeks or 16-weeks. (E) Intestinal permeability was tested by oral administration of FITC-dextran at 16th week of ingestion period. The serum FITC-dextran concentration appeared higher in CRD-animals, indicating that chronic CRD may disrupt intestinal barrier. (F, G) Intestinal transit ratio in charcoal meal test (F) together with the defecation frequency (the number of fecal pieces in 24 hours) and total stool weight (G) represents intestinal dysmotility in CRD-fed animals. (G) The length of large intestine did not differ between SD- and CRD-fed animals. Data are expressed as means±SEM of 5-11 mice/group. Statistical significant is indicated by \**p* < 0.05, \*\**p* < 0.01, \*\*\**p* < 0.005, \*\*\*\**p* < 0.001 according to unpaired *t-test* or Mann Whitney test.

One of the most known roles of intestinal SCFA is the maintenance of intestinal barrier and motility(26,27). We thus focused on the physiological changes in the intestinal tract and evaluated the intestinal permeability and motility. FITC-dextran test revealed that chronic CRD consumption will induce the intestinal hyperpermeability (Figure 4E). CRD consumption also induced intestinal dysmotility, resulting in the decrease of defecation frequently and stool weight (Figure 4F, G). However, chronic CRD had no significant effect to expression of gene related to the intestinal barrier (Figure S5). In addition, we confirmed the presence of intestinal inflammation by colorectal shortening and cytokine expression and found no significant differences in the length of large intestine (Figure 4H) and the mRNA expression of cytokines (Figure S6). Although no severe molecular difference or inflammation has been observed, these results indicate that CRD may induce the deterioration of intestinal environment including dysbiosis (decreased SCFA), dysmotility of the intestine and disruption of intestinal barrier.

### CRD causes intestinal hypersensitivity by upregulation of TRPA1

We then assessed the intestinal responsiveness to noxious stimuli to investigate the possible effects of the biochemical and physiological changes in CRD-fed mice. Notably, CRD-fed mice displayed clear hypersensitivity to nociceptive stimuli by allyl Isothiocyanate (AITC, Figure 5A) and capsaicin (Figure 5B). We further found upregulation of intestinal transient receptor potential ankyrin 1 (TRPA1) (Figure 5E) and sodium glucose co-transporter1 (SGLT1) in CRD-fed group (Figure 5D), while TRP vanilloid 1 (TRPV1) expression had no change (Figure S7B). AITC is the activator of TRPA1, and SGLT1 is known to fire the vagus nerve together with TRPA1 and TRPV1(28). These results therefore indicate that CRD consumption lead the upregulation of TRPA1 and SGLT1, which may enhance the responsiveness to noxious stimuli resulted as the intestinal hypersensitivity and further vagus nerve firing.

**Figure 5:**
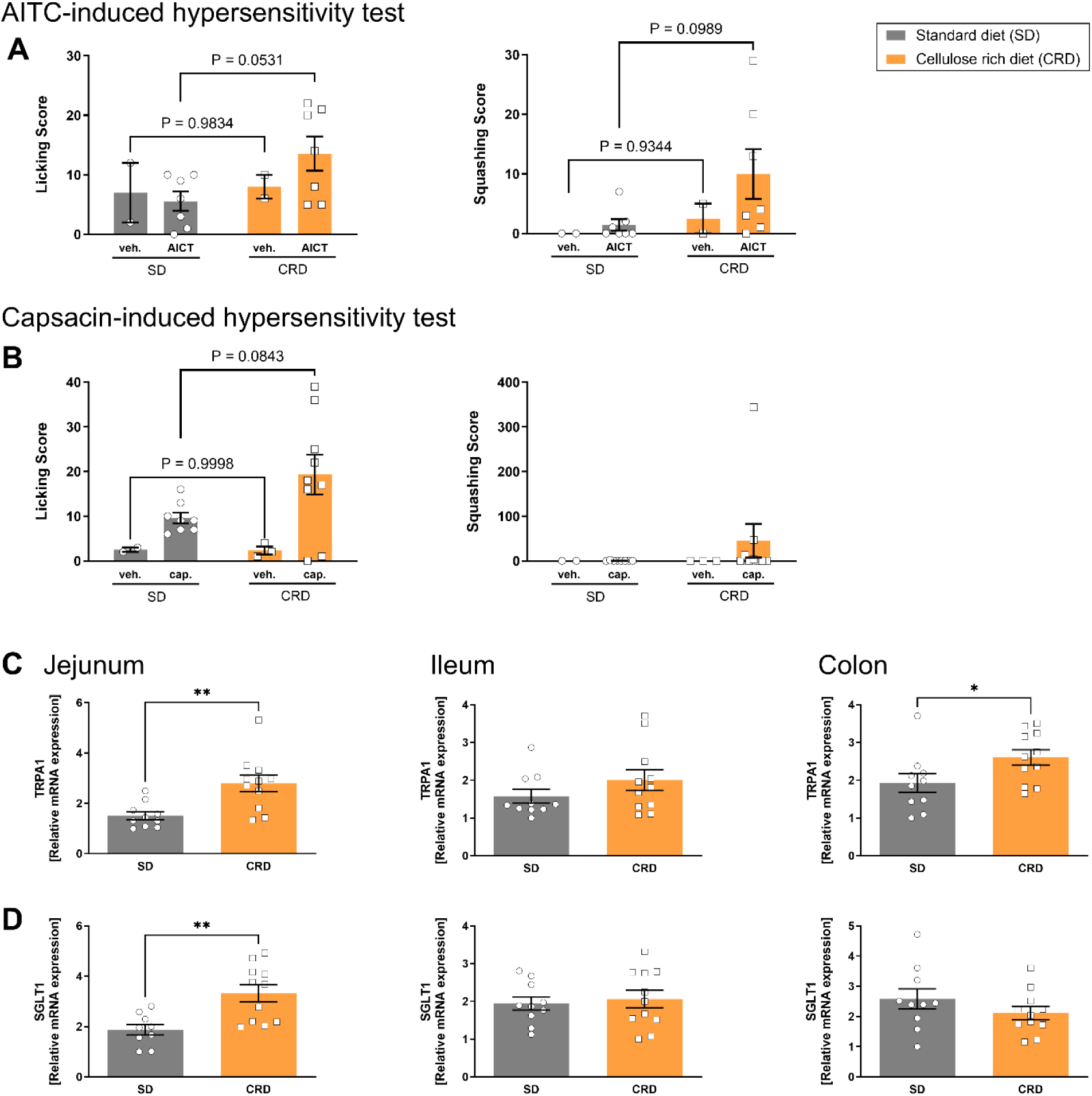
Chronic CRD-exposure evoked the intestinal hypersensitivity possibly through increased TRPA1. (A-B) Hypersensitivity was examined using the intestinal stimulation by ctcelomically administered AITC (A) and capsaicin (B) at 16th week of ingestion period. Licking of the abdomen and squashing of lower abdomen against the floor was taken as the pain-related behavioral score. CRD-exposed animals displayed increased hypersensitivity in both tests. (C-D) The mRNA expression level of TRPA1 (C) and SGLT1 (D) was increased in intestine tissues of mice already after 8 weeks of CRD exposure. (E-F) Effect of opioidergic system inhibition on anxiety-like behavior (E) and locomotive activity (F) was assessed animals after 16-week exposure of SD or CRD. Anxiety-like behavior was measured by marble burying test. Locomotive activity was assessed by open field test immediately after marble burying test. Pretreatment of naloxone (1mg kg^-1^) could suppress the CRD-evoked anxiety with no effect to the locomotive activity. Data are expressed as means±SEM of 5-11 mice/group. Statistical significant is indicated by \**p* < 0.05, \*\**p* < 0.01 according to two-way ANOVA with Sidak’s multiple comparisons test, unpaired *t-test*, or Mann Whitney test.

### CRD-induced intestinal hypersensitivity may enhance the anxiety and the amygdalar dopamine release through opioidergic systems

Recent study showed that gut-to-brain vagal transmission will activate the endogenous opioidergic systems in brain(29). It is also known that noxious stimuli or chronic hypersensitivity will activate opioidergic systems by tonic release of endogenous opioids such as enkephalin(30). Thus, we hypothesized that intestinal hypersensitivity evoked by chronic CRD may also cause the activation of the opioidergic systems, leading to over-activation of the amygdala dopaminergic system. To determine the contribution of opioid receptors, we injected the opioid receptor antagonist naloxone to mice and examined whether the CRD-induced anxiety could be inhibited by the blockage of opioid signaling. As expected, naloxone injection suppressed the CRD-induced anxiety (Figure 6B) with no effect to the locomotor activity (Figure 6C). These results suggests that CRD consumption evoked intestinal hypersensitivity to lead the excessive vagus nerve firing, which may activate the opioidergic system to increase the dopamine release in amygdala, that in the end lead to the enhanced anxiety.

**Figure 6:**
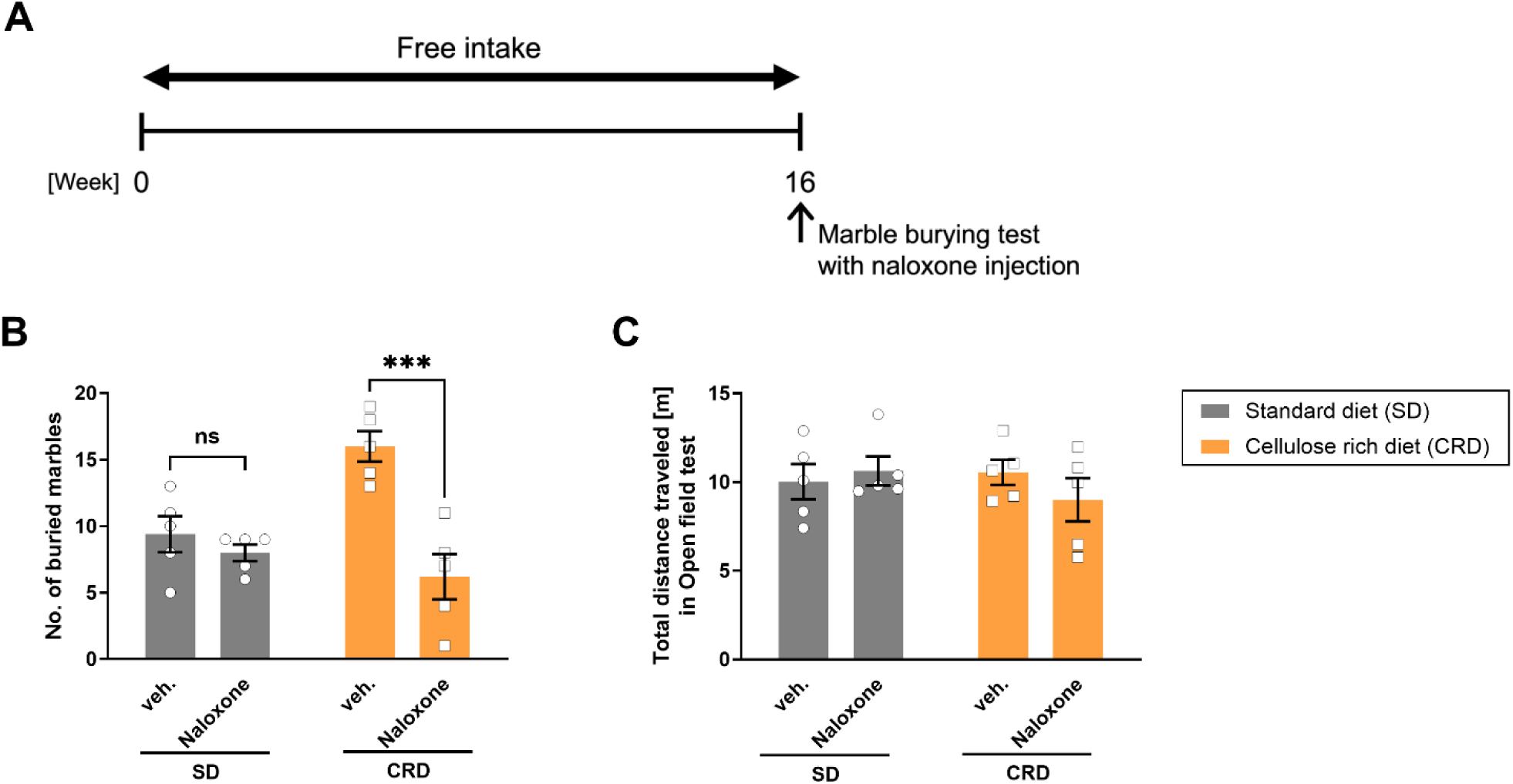
CRD-induced anxiety has been suppressed by opioid receptor antagonist. (A) Overall scheme of the experiments. The marble burying test with naloxone injection was conducted after 16 weeks of CRD exposure. (B-C) Effect of opioidergic system inhibition on anxiety-like behavior (B) and locomotive activity (C) was assessed animals after 16-week exposure of SD or CRD. Anxiety-like behavior was measured by marble burying test. Locomotive activity was assessed by open field test immediately after marble burying test. Pretreatment of naloxone (1mg kg^-1^) could suppress the CRD-evoked anxiety with no effect to the locomotive activity. Data are expressed as means±SEM of 5 mice/group. Statistical significant is indicated by \*\*\**p* < 0.005 according to two-way ANOVA with Sidak’s multiple comparisons test.

## Discussion

In the present study, long-term ingestion of CRD induced the chemical and physiological intestinal change, such as decreased SCFA, intestinal hypersensitivity, and increased intestinal permeability. We also found that chronic CRD evoked overactivation of amygadalar dopaminergic system via vagus nerve then opioidergic system, which resulted as enhanced anxiety possibly mediated through brain-gut axis.

We could observe CRD-induced anxiety particularly in the marble burying test, but not in the open field test and the elevated plus-maze test. This could be explained by the neuronal mechanisms underlying respective behavioral tests used to assess anxiety. While repeated administration of serotonin reuptake inhibitors can suppress marble burying behavior in mice(31), it is ineffective for anxiety-like behaviors in the open field test and the elevated plus maze test(32). Our finding suggests that CRD-induced anxiety which is only found in marble burying test may be regulated through different neuronal mechanisms e.g. serotonoergic systems, which requires more detailed assessments.

We also found significant increase of serum cortisol concentration in CRD group, accompanied with enhanced anxiety. Interestingly, the urinary cortisol concentration was significantly decreased in anxiety-evoking CRD-fed mice. There are two possible reasons for this urinary corticosterone reduction: The first is the corticosterone dysfunction by chronic physiological stress. Repeated surges of cortisol by chronic stress lead cortisol dysfunction, like depletion of cortisol or hypersensitivity of the negative feedback system(33). Salivary cortisol levels are decreased in patients with chronic lumbago(34). Further, HPA axis negative feedback was enhanced by dexamethasone administration in chronic myogenous facial pain patients (35). Based on these findings, it is possible that e.g. chronic pain or hypersensitivity enhances negative feedback of the HPA axis and suppresses corticosterone secretion. The second possible reason is excessive opioidergic system activation by intestinal hypersensitivity. As opioid agonists decrease urinary corticosterone levels(36), CRD-induced overactivation of the opioid system may suppressed homeostatic corticosterone secretion. Despite of various possible mechanisms, we expect that this corticosterone reduction might be a cause of CRD-induced intestinal dysmotility, as corticosterone secretion has been suggested to cause increased peristalsis(37).

Chronic CRD exposure not only induced anxiety but also lead alteration of dopamine levels together with decrease of D2Rs in amygdala. Amygdala is known for its regulatory role for anxiety and fear, which could be demonstrated by excessive activation of amygdala in patients of psychiatric disorders with symptoms of excessive anxiety, such as depression, anxiety disorders, and PTSD(38). Notably, microinjection of D1R and D2R agonists or antagonists into the amygdala produce anxiogenic or anxiolytic effects, respectively(39). Moreover, chronic dopamine agonists exposure decreases D2R levels on dopaminergic neurons(40). Altogether, increase of amygdalar dopamine level evoked by CRD exposure might lead D2R reduction, which may modify the dopaminergic system resulting to anxiety-like behavior as the outcome.

In addition to brain monoamine, recent study suggested that the brain inflammation may also involve in emotion. Thus, the increase of inflammatory cytokines IL-6 and TNF-α in the amygdala is associated with the induction of anxiety-like behavior(23), and suppression of amygdalar TNF-α ameliorates anxiety-like behavior(41). We therefore confirmed whether anxiety-evoking chronic CRD exposure caused the inflammation in the brain. Surprisingly, CRD decreased the mRNA expression of IL-6 and TNF-α in the amygdala. Interestingly, decrease of CD11b positive monocyte population in the bone marrow and spleen is also found in CRD-fed mice. Monocytes have been reported to circulate through blood vessels and release TNF-α in the brain, involving the synaptic turnover(42). Thus, reduced amygdalar TNF-α may be due to the decreased monocyte level in the bone marrow and spleen by long-term CRD exposure.

Our study clarified that the worsening of intestinal environment as the major cause of neuronal change and following enhanced anxiety. Long-term CRD evoked decreased SCFA concentrations which caused increase of cecum pH, reflecting the deterioration of intestinal environment. CRD consumption also increased intestinal permeability along with decrease of intestinal motility and defecation frequency, suggesting the global exacerbation of intestinal condition. It is known that SCFA promotes serotonin production in the intestinal tract to maintain normal peristalsis(43). Butyrate, one of the SCFAs, increase ChAT immunoreactive neurons in intestinal tissue, enhancing colonic contractions due to cholinergic innervation(27). SCFAs also have a role in enhancing tight junctions and maintaining intestinal barrier(26), which is often lost in several psychiatric diseases(44). Interestingly, vagotomized mice showed no anxiety or amygdalar dopamine increase, despite of gut condition aggravation like decreased SCFA as well as increased cecum pH. All in all, SCFA downregulation by CRD exposure may cause adverse physiological effects on the gut e.g. increased intestinal permeability and dysmotility, leading the negative psychiatric effect through the gut-brain axis mediated by vagus nerve.

Another interesting finding on effect of CRD is the intestinal hypersensitivity to the noxious stimuli given by capsaicin or AITC. We also found the particular increase on TRPA1, one of the most known pain-related cation channels, which explains the revelation of intestinal hypersensitivity by CRD. In general, TRP family proteins are receptors that respond to temperature, pH, osmotic pressure, chemical stimuli, and mechanical stimuli in peripheral nerves and transmit these stimuli to the central nervous system by activating nociceptors(45). TRPA1 has been shown to respond strongly to mechanical and chemical stimuli such as AITC. TRPA1 is also characteristically expressed in enterochromaffin cells (ECs) which activate the vagus nerve via serotonin secretion(46). These findings suggest that CRD-induced increased expression of TRPA1 induces intestinal hypersensitivity, evoking excessive vagal signaling.

There are three possible reasons for the increased expression of TRPA1. First, it is suggested that TRPA1 activation promotes serotonin production in ECs, which in turn stimulates peristalsis(46). As CRD exposure caused decreased peristalsis, TRPA1 might be upregulated to restore the peristalsis to the normal state. Second, it is known that inflammation causes the increase of TRPA1 and number of TRPV1/TRPA1-responsive neurons in a mouse model of caerulein-induced pancreatitis(47), suggesting that inflammation regulates TRPA1 expression and activity. As ileum IL-1βtended to increase by CRD exposure, that slight enhancement of inflammation may have triggered the increase of TRPA1. Finally, past study showed the contribution of reactive oxygen species to TRPA1 upregulation through NRF2 activation(48), which could be decreased by SCFA(49). Therefore, the decrease in SCFA concentrations due to CRD consumption may increase oxidative stress, which may upregulate the TRPA1 expression by NRF2 activity. As CRD consumption also upregulated SGLT1, which is reported to be increased in H_2_O_2_-treated ECs(50), we expect that contribution of oxidative stress might be the most plausible theory.

Past reports demonstrated that TRPA1 and SGLT1 might stimulate vagus signaling(28,47), which may link to the dopaminergic system in brain. The vagus nerve is known to project to nucleus tractus solitarii, and suggested to affect amygdalar activity indirectly via other brain region, such as ventral tegmental area and locus ceruleus(51,52), possibly through activation of enkephalin-mediated endogenous opioidergic systems(29). We could confirm the contribution of the endogenous opioid system to CRD-evoked anxiety-like behavior by acute naloxone administration. Endogenous opioid system indirectly promotes dopaminergic activation at amygdala through inhibition of the GABAergic system in the ventral tegmental area(53). We therefore propose the following hypothetic scheme: CRD-induced TRPA1 upregulation first causes increased transmission of nociceptive stimuli to the brain through vagus nerve, which may activate the endogenous opioid system which lead activation of dopaminergic neurons at VTA, which could increase dopamine release at the amygdala in CRD-fed mice (shown in the Graphical Abstract). However, the dopamine level in the nucleus accumbens were not increased in CRD-fed mice (Figure S2C).

We also can not exclude the possible contribution of intestinal microbiota on the present findings. Past study showed that chronic consumption of AIN-93G modified gut microbiota in mice(54). Thus, AIN-93G increased presence of Allobaculum to be the major genus (43.4 %) of mice. This result suggests two important aspects: 1) AIN-93G lead significant loss of microbiome diversity, and 2) AIN-93G increased particular microbiota Allobaculum, which is frequently observed as the characteristic of aged mice. Since CRD (AIN-93M) has similar composition as AIN-93G, it is possible that our model exhibit similar changes in the gut microbiota and affect gut-brain axis to modify emotion. Altogether, other molecular mechanisms or pathways may exist to lead CRD-evoked anxiety, which requires further detailed examination.

In summary, our novel findings suggest that CRD consumption cause 1) the intestinal deterioration and hypersensitivity, which leads 2) abnormality in amygdalar dopaminergic system mediated by vagus nerve then opioidergic system, resulting as 3) enhanced anxiety. Number of recent evidence supports that dietary fiber consumption is beneficial to health by reducing the risk of various diseases such as metabolic disorders, cardiovascular diseases, and colorectal cancer(55). Although the proportions of each nutrient are different in MF and AIN-93M (Table 1-3), there is no doubt that dietary fiber has the greatest effect on the intestine(19). Present study shows the first time, the particular kind of dietary fiber, in accurate the insoluble dietary fiber, may cause emotional problems through the modification of gut environment. Our study confirms that diet affects the risk of developing psychiatric disorders, suggesting the importance of further detailed examination of diets to clarify e.g. whether particular food/nutrition has anxiogenic or anxiolytic effect, which in the end might enable the prevention and treatment of psychiatric disorders through dietary intervention.

**Table 1.**
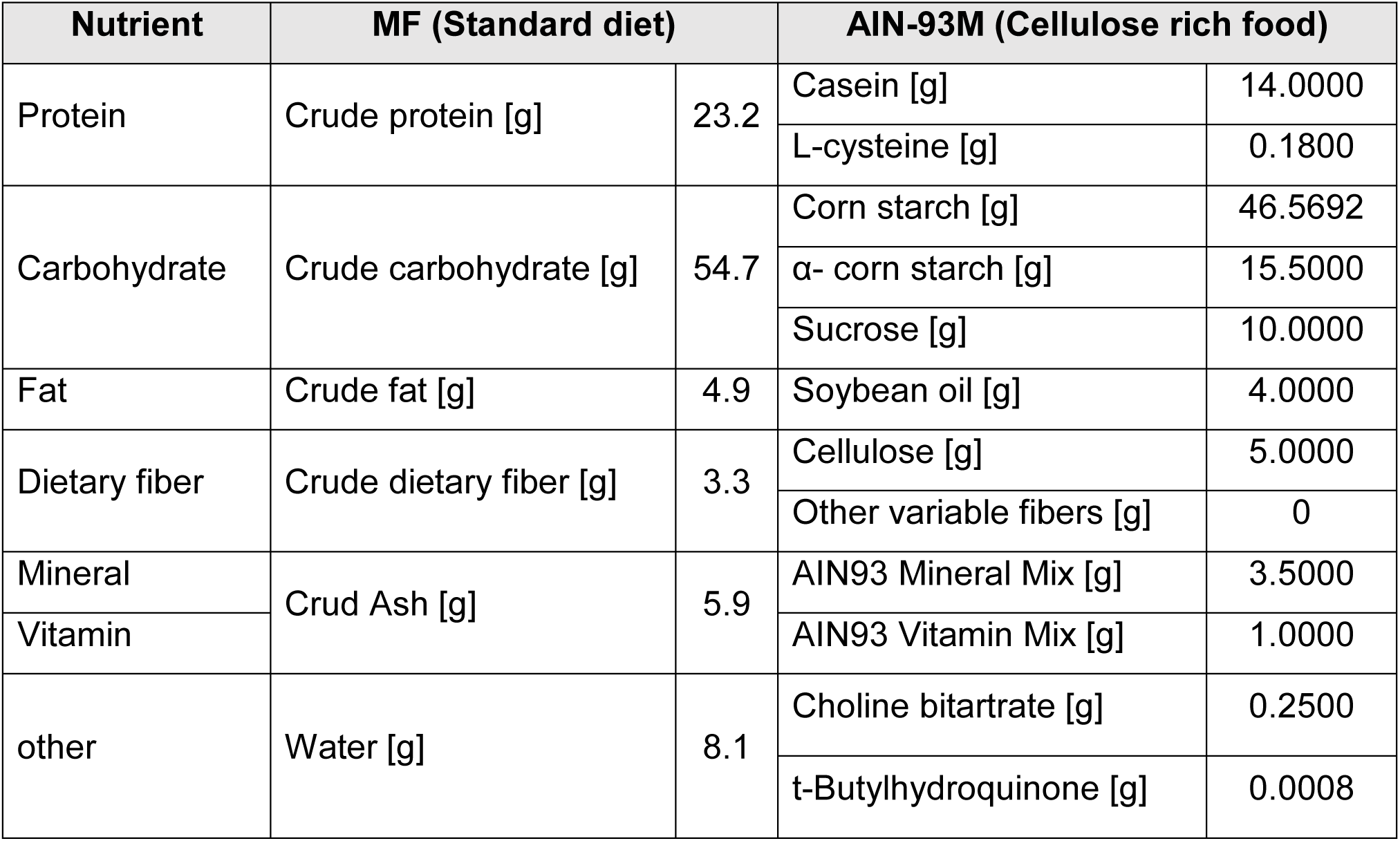
Nutritional composition of the diets [%]. Created with reference to the website of Oriental Yeast Co., Ltd. and CLEA Japan, Inc. (MF: https://www.oyc.co.jp/bio/LAD-equipment/LAD/ingredient.html, AIN-93M: https://www.clea-japan.com/company/outline.html) *Detailed composition is described in Table 2. ** Detailed composition is described in Table 3.

**Table 2.** Detailed nutrient composition of MF (Amount contained in 100 g) Created with reference to the website of Oriental Yeast Co. (https://www.oyc.co.jp/bio/LAD-equipment/LAD/ingredient.html).

| Mineral |  | Vitamin |  | Amino acid |  |
| --- | --- | --- | --- | --- | --- |
| Calcium [g] | 1.04 | Vitamin A* [IU] | 1638 | Isoleucine [g] | 0.92 |
| Phosphorus [g] | 0.81 | Vitamin D3 [IU] | 111 | Leucine [g] | 1.77 |
| Magnesium [g] | 0.24 | Vitamin E [mg] | 8.9 | Lysine [g] | 1.27 |
| Natrium [g] | 0.21 | Vitamin K3** [mg] | 0.04 | Methionine [g] | 0.43 |
| Kalium [g] | 0.99 | Vitamin B1 [mg] | 1.92 | Cystine [g] | 0.36 |
| Iron [mg] | 11.1 | Vitamin B2 [mg] | 1.01 | Phenylalanine [g] | 1.06 |
| Copper [mg] | 0.74 | Vitamin C [mg] | 5 | Tyrosine [g] | 0.74 |
| Zinc [mg] | 5 | Vitamin B6 [mg] | 0.87 | Threonine [g] | 0.89 |
| Manganese [mg] | 5.2 | Vitamin B12 [μg] | 4.6 | Tryptophan [g] | 0.3 |
|  |  | Inositol [mg] | 467.3 | Valine [g] | 1.11 |
|  |  | Biotin [μg] | 30.1 | Arginine[g] | 1.47 |
|  |  | Pantothenic acid [mg] | 2.14 | Histidine [g] | 0.62 |
|  |  | Niacin [mg] | 10.22 | Alanine [g] | 1.19 |
|  |  | Choline [g] | 0.18 | Aspartic acid [g] | 2.12 |
|  |  | Folic acid [mg] | 0.16 | Glutamic acid [g] | 3.94 |
|  |  |  |  | Glycine [g] | 1.15 |
|  |  |  |  | Proline [g] | 1.28 |
|  |  |  |  | Serine [g] | 1.1 |

**Table 3.** Detailed nutrient composition of AIN-93M [%]. Created with reference to the website of CLEA Japan, Inc. (https://www.clea-japan.com/company/outline.html).

| AIN93 Mineral Mix |  | AIN93 Vitamin Mix |  |
| --- | --- | --- | --- |
| CaCO<sub>3</sub> | 35.7000 | Sucrose | 97.3474 |
| Sucrose | 20.9782 | Vitamin E (50 %) | 1.5000 |
| KH<sub>2</sub>PO<sub>4</sub> | 25.0000 | Nicotinic acid | 0.3000 |
| NaCl | 7.4000 | D-Pantothenic acid Ca | 0.1600 |
| K<sub>3</sub>C<sub>6</sub>H<sub>5</sub>O<sub>7</sub>·H<sub>2</sub>O | 4.6600 | D-Biotin (100 %) | 0.0020 |
| K<sub>2</sub>SO<sub>4</sub> | 2.8000 | Vitamin B2 (Over 98 %) | 0.0600 |
| MgO | 2.4000 | Vitamin B6 | 0.0700 |
| FeC<sub>6</sub>H<sub>5</sub>O<sub>7</sub>·XH<sub>2</sub>O | 0.6060 | Vitamin B1 | 0.0600 |
| ZnCO<sub>3</sub> | 0.1650 | Vitamin A (325,000 IU/g) | 0.1231 |
| Na<sub>2</sub>SiO<sub>3</sub>·9H<sub>2</sub>O | 0.1450 | Vitamin D3 (100,000 IU/g) | 0.1000 |
| MnCO<sub>3</sub> | 0.0630 | Folic acid | 0.0200 |
| CuCO<sub>3</sub>Cu (OH)<sub>2</sub> | 0.0324 | Vitamin B12 (0.1 %) | 0.2500 |
| CrK (SO<sub>4</sub>)<sub>2</sub>·12H<sub>2</sub>O | 0.0275 | Vitamin K1 | 0.0075 |
| H<sub>3</sub>BO<sub>3</sub> | 0.0082 |  |  |
| NaF | 0.0064 |  |  |
| NiCO<sub>3</sub>·2Ni (OH)<sub>2</sub>·4H<sub>2</sub>O | 0.0032 |  |  |
| LiCl | 0.0017 |  |  |
| Na<sub>2</sub>SeO<sub>4</sub> | 0.0010 |  |  |
| KIO<sub>3</sub> | 0.0010 |  |  |
| (NH<sub>4</sub>)<sub>6</sub>Mo<sub>7</sub>O<sub>24</sub>·4H<sub>2</sub>O | 0.0008 |  |  |
| NaVO<sub>3</sub> | 0.0007 |  |  |

## Methods and Materials

### Animals

Male ICR mice (Tokyo Laboratory Animals, Japan) aging 8-weeks-old at the beginning of the experiments were used. All mice were kept at a room maintained at 22±2°C, humidity of 60±5%, with a 12-h light (100-150 lux): 12-h dark cycle (light on between 8:00 and 20:00). Food and water were available ad libitum except during behavioral observations. Total of two cohorts are prepared: MF (Oriental Yeast Co., Ltd., Tokyo, Japan)-fed standard diet (SD) group and AIN-93M (CLEA Japan, Inc., Tokyo, Japan)-fed cellulose rich food (CRD) group. We defined AIN-93M as a “high cellulose diet” because AIN-93M contains only 5% cellulose as dietary fiber, while MF contains 3.3% crude fiber, both soluble and insoluble. Table 1-3 shows the detailed composition of them. The duration of the dietary intervention was up to 18 weeks, with major behavioral tests and dissection at weeks 8 and 16. Experiments were conducted in accordance with permission from the Committee for Animal Experimentation of the School of Science and Engineering at Waseda University (permission #A23-093), and in accordance with the Law (No. 105) and Notification (No. 6) of the Japanese Government.

### Drugs

Most of reagents and drugs has been purchased from Fujifilm Wako Pure Chemicals, Osaka, Japan, and antibodies has been purchased from Beckton Dickinson, NJ, USA, otherwise stated in the methods.

### Behavioral tests

Behavioral tests to assess anxiety (marble burying test, open field test, elevated plus maze) and intestinal hypersensitivity (AITC- or Capsaicin-induced hypersensitivity) were conducted. Respective test has been done at 2-4th, 6-8th and 16-18th week of feeding period with one-week interval for each testing. All experiments were conducted 17:00-20:00 due to avoid changes in behavior due to time of day, and data was assessed by automated data collection with software or individual observer by blind manner.

### Marble burying test

Marble burying test was conducted with slight modifications to previous studies(56) at 4, 8, and 16th week of feeding period. Individual mice were placed in a cage (24 x 37 x 17 cm) containing 20 marbles of various colors with a diameter of 1.5 cm arranged in a 4 x 5 grid, each on top of 5-cm of white softwood bedding, then left undisturbed for 15 minutes. A marble with more than two-third of its height covered with bedding was considered buried marbles. For each test cage, digital photographs were obtained at uniform angles and distances and buried marbles are counted by independent observer.

### Open Field Test

Open filed test was conducted at 2, 6, and 17th week of feeding period. Mouse was individually placed at the corner of a cage (24 x 37 x 17 cm) and allowed to move freely for 5 minutes. The locus of its activity was recorded by video camera and analyzed using a video tracking software ANY-maze (Stoelting, USA). The number of center-zone entry, center-zone staying time, traveled distance was measured and assessed.

### Elevated Plus Maze Test

Elevated plus maze test was conducted at 3, 7, and 18th week of feeding period. The plus maze was consisted of two 29 x 7 cm opened arms, two 29 x 7 cm closed arms, and a 7 cm square center zone. Mouse was placed at its center and allowed to move freely for 5 minutes. Its activity locus was recorded by video camera and analyzed using a video tracking software ANY-maze (Stoelting, USA). The staying time in the open and closed arm, the distance travelled in the open and closed arm, and the number of entries in the open and closed arm were measured and assessed.

### AITC-induced hypersensitivity

Allyl isothiocyanate (AITC)-induced hypersensitivity was assessed using the methods described previously(57). Mice were habituated in the 15 cm×10 cm×8 cm observation cage 20 minutes. AITC (2% vol vol^-1^ in rape seed oil; Nacalai Tesque Inc., Kyoto, Japan) was administered in 0.05 mL intrarectally and their behavior was then observed for 15 min. Pain-related behaviors were determined as follows; licking of the abdomen, stretching the abdomen, squashing of lower abdomen against the floor, and immobility. Behaviors were assessed in 5-second intervals and scores were calculated as follows: 2 points for pain-related behavior for at least 3 out of 5 seconds, 1 point for less than 3 out of 5 seconds, and 0 points for never performing the behavior during the 5 seconds. Immobility was measured from the point at which it stopped for more than 5 seconds.

### Capsaicin-induced hypersensitivity

Capsaicin-induced hypersensitivity was assessed using the methods described previously(58). Mice were habituated in the 15 cm×10 cm×8 cm observation cage for 20 minutes. Capsaicin (0.1% wt vol^-1^ in 4% Tween80/0.9% saline) was administered in 0.1 mL intrarectally and their behavior was then observed for 15 min. Pain-related behaviors were assessed and scored as described above in the AITC-induced hypersensitivity test.

### Naloxone administration

Naloxone (1 mg kg^-1^) were dissolved in saline and injected i.p. to mice 20 minutes prior to the marble burying test. The test was conducted by crossover trial with 4 days of recovery period. Effect of naloxone administration to the locomotor activity was assessed by open field test immediately after the marble burying test.

### Tissue collection

Tissue collection has been done 3-5 days after the final marble burying tests. Briefly, mice were euthanized by rapid decapitation, then brain, spleen, bone and whole gut (between pylorus to rectum) has been quickly harvested on ice. Then brain has been separated into prefrontal cortex (PFC), amygdala, striatum, nucleus accumbens and hippocampus for further use on monoamine measurement and RT-PCR. Spleen and bone were processed for flow cytometry. Gut has been separated into cecum, jejunum, ileum and colon for further use on short-chain fatty acid measurement and RT-PCR.

### Brain monoamine levels Measurement

Brain monoamine levels were measured by high performance liquid chromatography (HPLC) as previously reported(59). Samples were first processed with 0.2 M perchloric acid solution (containing 100 M EDTA-2Na) together with isoproterenol (Sigma-Aldrich, USA) as an internal standard, followed by ultrasonic homogenizing and centrifugation at 15,000 rpm for 10 minutes to collect supernatant containing monoamines. Samples and the standards were then purified through a 0.45 µm filter (EMD Millipore, USA) and injected into an HPLC coupled with electrochemical detector (HTEC-510, Eicom Co., Kyoto, Japan). Monoamine levels of samples were measured based on the measurement result of standard. The mobile phase was 0.1 M acetate-citrate buffer (containing 5 mg L^-1^ EDTA·2 Na, 190 mg L^-1^ 1-octanesulfonic acid sodium salt, and 15 % methanol). The flow rate was 500 μl min^-1^, the applied column temperature was 25°C, and the voltage was +750 mV for Ag/AgCl, respectively. For data analysis, EPC-300 software (version 2.5.10, Eicom) was used.

### RNA Extraction

mRNA expression levels were measured by real-time RT-PCR, using the described protocol from previous study(41). The brain tissue was collected directly in a tube containing 500 µL of Trizol (Ambion, USA) then homogenized using Tissuelyser II (Qiagen, Germany). Then 500 µL of chloroform was added and centrifuged at 15,000 rpm for 10 minutes to collect supernatant. The one-third supernatant volume of CIA solution (chloroform: isoamyl alcohol = 49:1) was added and further centrifuged at 11,500 rpm for 10 minutes. The supernatant was then transferred to another tube, 100 µL of 3 M sodium acetate and 100 µL of isopropanol were added, and incubated at room temperature for 20 minutes. Sample was centrifugated at 11,500 rpm for 20 minutes, followed by pellet washing by 80 % ethanol, then dissolved into 20 µL of DEPC (NIPPON GENE Co., Ltd., Tokyo, Japan) for further real-time RT-PCR.

### Real-time RT-PCR

The RNA samples were diluted with DEPC to achieve a concentration of 50 mg/mL of RNAs using a spectrophotometer (NanoDrop, Thermo Fisher Scientific, USA). The adjusted samples were subjected to real-time RT-PCR using the One Step SYBR RT-PCR Kit (Takara Corporation, Tokyo, Japan) and the PIKO REAL 96 Real-Time PCR System (Thermo Fisher Scientific, USA). The primer sequences used to amplify each gene and the RT-PCR settings are shown in Supplementary Tables 1 and 2. The relative expression of each target genes was normalized to that of 18s rRNA, and the data were analyzed by the ΔΔCt method.

### Vagotomy

Vagotomy was conducted with slight modifications to previous study(60). Vagotomy was performed under combined triadic anesthesia (Medetomidine Hydrochloride (domitor, ZENOAQ, Japan), Butorphanol Tartrate (vetorphale, Meiji, Japan), midazolam (Sandoz Corporation, Japan)). A laparotomy incision was made on the ventral midline and the abdominal muscle wall was opened with a second incision. The gastrohepatic ligament was severed using fine forceps, and the stomach was gently retracted, revealing the descending ventral oesophagus and the ventral subdiaphragmatic vagal trunk. The hepatic branch of this vagal trunk was then transected using fine forceps. Sham group was produced with same procedure except vagal trunk transection. Successful vagotomy was confirmed by response to CCK-8 (8 μg kg^-1^, i.p.). Mice with lower food intake than the average of CCK-8-treated control group were considered as false-vagotomized, and removed from experimental group. The results of CCK test are shown in Supplementary Figure 8 and Supplementary Table 4.

### Cecal pH and short-chain fatty acid (SCFA) Measurement

For cecal pH measurement, the probe of pH meter (Euthech Instruments, USA) was inserted directly into a dissected cecum, then waited until the value has been stopped fluctuating. Then cecum content was rapidly collected on ice, then processed for gas chromatography coupled with mass spectrometry (GC-MS) to measure the cecum SCFA levels, using the protocol previously described(61), using model 7890B or 5977B instruments (Agilent Technologies, Inc., Santa Clara, CA, USA). 50 µL of sulfuric acid, 200 µL of chloroform and diethyl ether were added to approximately 50 mg of cecum contents. Then 100 µL of trimethylsilylation reagent (TMSI-H; GL Science Inc., Tokyo, Japan) was added to 300 µL of supernatant containing SCFAs to convert target compounds into volatile and thermally stable derivatives. The sample was then incubated at 60°C for 30 minutes and on ice for 10 minutes. It was centrifuged at 14,000 rpm at room temperature for 30 seconds and 1 µL of its supernatant was subjected to GC-MS. To generate the calibration curve, a standard mixture containing acetic acid, propionic acid, lactic acid, and butyric acid was also subjected to GC-MS after operating as in the sample. The capillary column was InertCap Pure-WAX (30 m × 0.25 mm, df = 0.5 µm) (GL Sciences, Japan), and helium gas was used as the carrier gas during the measurement to increase the initial temperature from 80°C to a final temperature of 200°C.

### Charcoal meal test

Charcoal meal test was conducted with slight modifications from previous study(58). Mice were fasted about 16 hours before the test. Vermilion Indian ink was orally administered to them at dose of 10 mL/kg body weight. Mice were sacrificed 10 minutes after administration and their duodenum and small intestine (between pylorus to the colon) were removed and spread on the aluminum plate immediately. Transit ratio was determined as the distance of ink transport divided by the total length of the intestinal tract.

### Intestinal permeability test

Intestinal permeability test was conducted with slight modifications from previous study(58). 200 μL of FITC-dextran (MW 4000) solution (50 mg/mL; Merck KGaA, Darmstadt, Germany) was orally administered to mice. Serum samples were collected from the orbital plexus under anesthesia with isoflurane, incubated at room temperature for 1 hours, centrifuged at 3,000 rpm for 20 minutes and the supernatant was collected. Following to the 5-fold sample dilution in the Milli-Q water, plasma FITC concentration was measured as fluorescence intensity using BioTek Synergy H1 (Agilent Technologies Japan, Ltd., Tokyo, Japan).

### Feces and Urine Collection

Mice were housed in the metabolic cage (Natsume Seisakusho Co., Ltd, Tokyo, Japan) for three-days then feces and urine samples are collected 24 hours at third day. Urine sample was centrifuged at 1,000 g and the supernatant was stored at −80°C until use to measure corticosterone levels. Defecation frequency was defined as the number of fecal pieces in 24 hours on the third day, and fecal mass was measured at that time.

### Corticosterone ELISA

Urine and Serum corticosterone levels were measured by using the ELISA kit (Funakoshi Co., Ltd., Tokyo, Japan). Corticosterone measurements were performed according to the instructions provided in the ELISA kit. In brief, urine sample was collected using metabolic cage as described above and diluted 20-fold in the buffer. Serum samples were collected from the orbital plexus under anesthesia with isoflurane, incubated at room temperature for 1 hours, centrifuged at 3,000 rpm for 20 minutes and the supernatant was stored at −80°C until use. Plasma samples were diluted 100-fold in the buffer and processed according to the instruction of the kit. The corticosterone concentration was measured as absorbance using BioTek Synergy H1 (Agilent Technologies Japan, Ltd., Tokyo, Japan).

### Flow cytometry

Spleen and bone marrow was used for the flow cytometry. Single cell suspensions were prepared from freshly harvested spleen and bone marrow using following procedure: First, spleens were mushed using a plunger and passed through a 70 μm filter with 0.5 mL 1xACK buffer. Bone marrow is flushed out from femur and tibia by 1mL FACS buffer. The cell suspension is hemolyzed with 1xACK buffer, then centrifuged at 1500 rpm for 10 min, and the supernatant was discarded. After another washing of cell suspension then further nonspecific Fc binding with anti-mouse CD16/32 (Bio X Cell, NH, USA), cell surface markers were stained with the following antibodies (Supplementary Table 3) : anti-mouse CD19-BV510 (GK1.5), CD5-BV421 (53-6.7), NK1.1-APC (PK136), CD11b-PerCP-Cy5.5 (M1/70), Ly6C-FITC (AL-21), Ly6G-eF450 (1A8, Thermo Fisher Scientific), I-A/I-E-APC-Cy7 (M5/114.15.2, Biolegend). Flow cytometry was performed on CytoFLEX S (Beckman Coulter) and analyzed using CytExpert software (Beckman Coulter). Gating strategy is shown in Figure S3C. Only the cell population displayed significant difference between the experimental cohort is chosen and analyzed.

### Statistical Analyses

All data were analyzed using GraphPad Prism version 6.03 (GraphPad Software, USA). First, we confirmed whether the data followed a normal distribution using the D’Agostino-Pearson test/Kolmogorov-Smirnov test, and whether the data were equal variances using the F value test/Bartlett’s test. Significance between two independent groups was assessed by parametric analysis using unpaired *t-tests* or by non-parametric analysis Mann-Whitney tests. Significance between three independent groups was assessed by parametric analysis using One-way ANOVA with uncorrected Fisher’s LSD, or One-way ANOVA with Tukey’s multiple comparisons test. Non-parametric analysis Kruskal-Wallis tests with Dunn multiple comparison tests were used for abnormally distributed data. Two-way ANOVA with Sidak’s multiple comparisons test was used for two factors data.

## Acknowledgements

We thank Dr. Daisuke Yamada and Dr. Hisamichi Yoshioka from Tokyo University of Science for technical supports and discussion. SS was supported by a Grant-in-Aid for Scientific Research (A, 19H01089) obtained from the Japan Society for the Promotion of Science, the JST-Mirai Program (JMPJM120D5). CN was supported by Home-Returning Researcher Development Research (19K24693) from the JSPS. AH was supported by a Grant-in-Aid for Young Scientists (19K14018) from the JSPS.

## Author Contribution

KI, CN and AH contributed to conception and design of the study with help of SS. KI mainly conducted the experiments with the help of HH for intestinal permeability test, flow cytometry, and its analysis, YK for charcoal meal, HS for GC-MS and HPLC, and AH for vagotomy and behavior tests. KI, AH, CN, and YK dissected and sampled the mice. All authors contributed to manuscript revision, read, and approved the submitted version.

