## Supplemental materials for "Cellulose Rich Food Leads Anxiety through Gut-Brain Axis-mediated Amygdalar Dopamine Upregulation"

### Supplemental Information

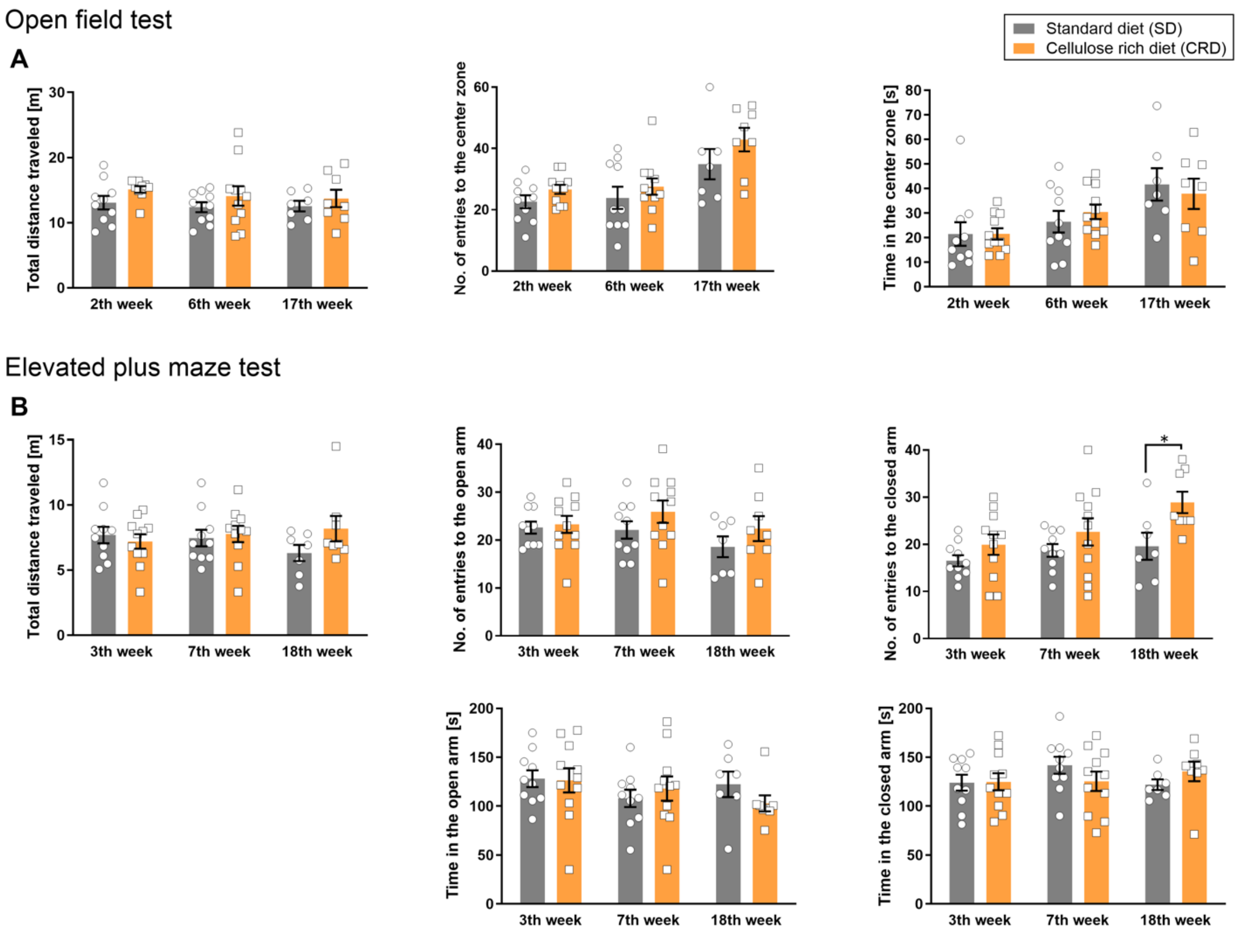


**Supplementary Figure 1: CRD consumption did not affect the anxiety-like behavior in open field test and elevated plus maze test.**

Anxiety behavior as well as locomotor activity was assessed using (A) open field test and (B) elevated plus maze test. Data are expressed as means±SEM of 9-11 mice/group. Statistical significance is indicated by *p < 0.05 according to Mann Whitney test.

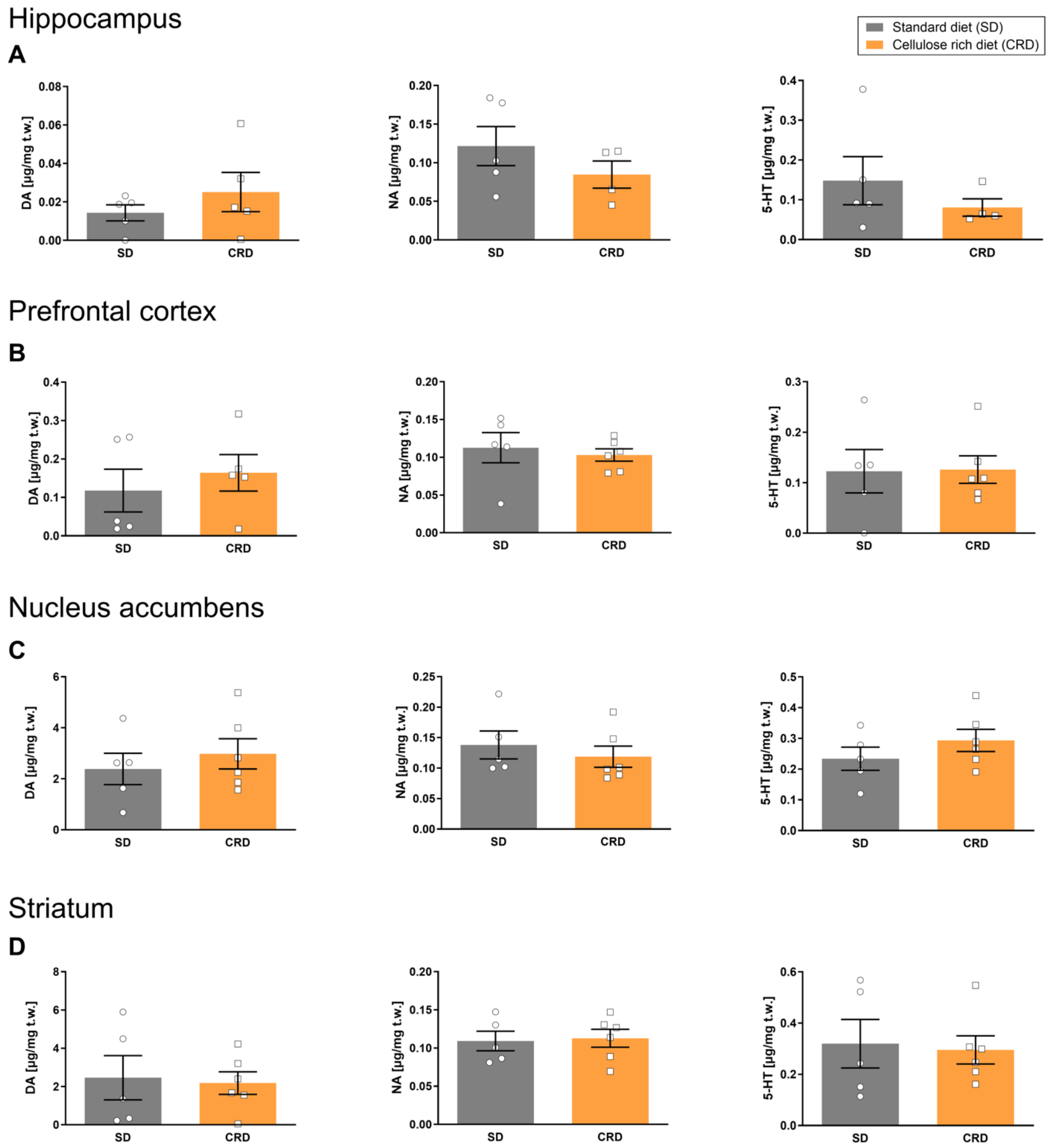


**Supplementary Figure 1: CRD did not affect any monoamine levels in other brain regions.**

The monoamine levels in the hippocampus (A), prefrontal cortex (B), nucleus accumbens (C) and striatum (D) were measured using brain sample from mice fed each diet for 16 weeks. Data are expressed as means±SEM of 5-8 mice/group.

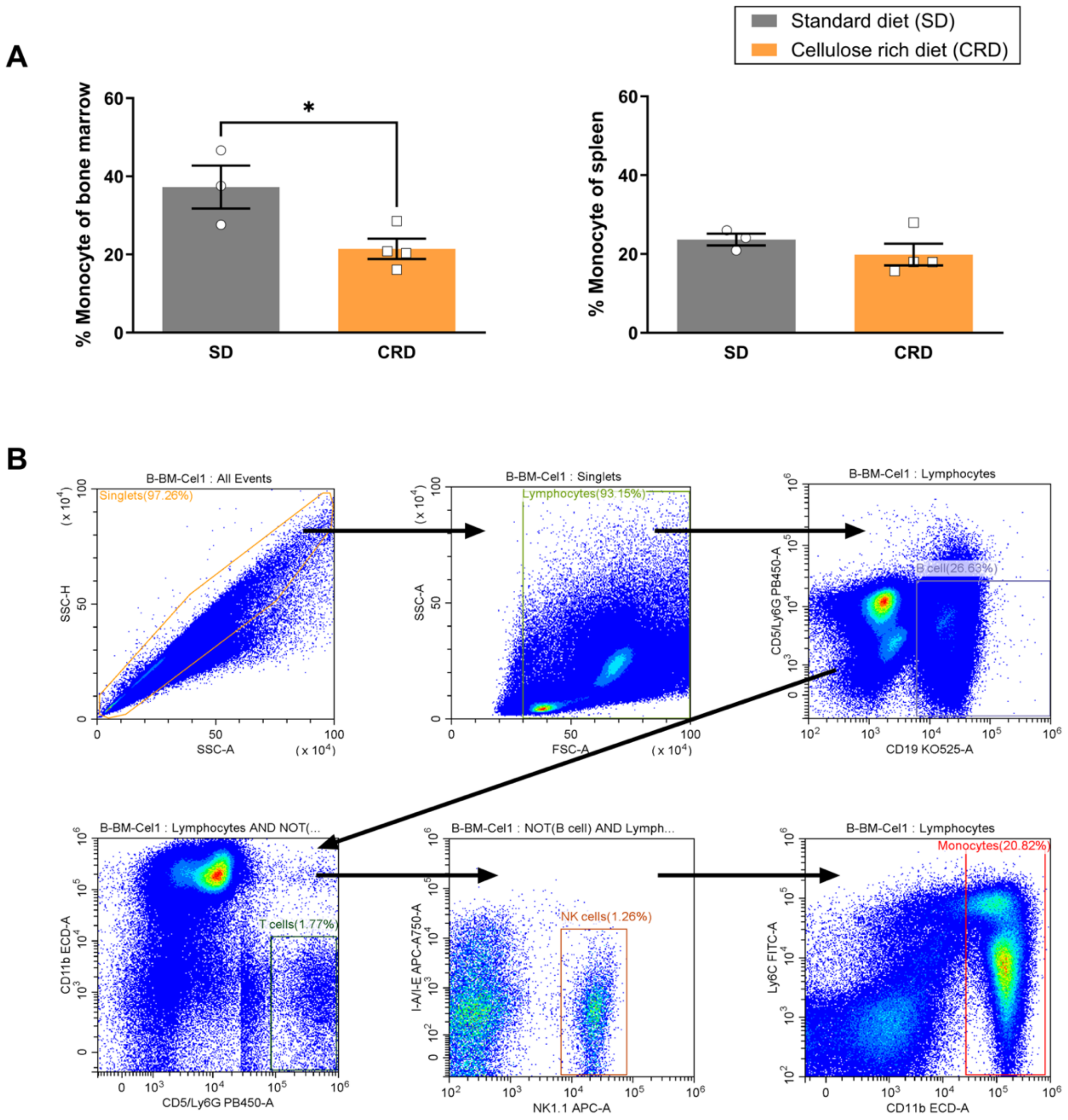


**Supplementary Figure 2: CRD exposure decreased the monocyte population in bone marrow.**

(A) The amygdalar mRNA expression level of cytokines was determined at 16th week after the exposure to either SD or CRD. (B) The monocyte population in bone marrow and splenocytes was compared between SD- or CRD-fed mice at feeding period of 16th week. (C) The gating strategy of present flow cytometric data. Monocyte population was defined as

CD5⁻CD19⁻NK1.1⁻CD11b⁺ cells. Data are expressed as means±SEM of 9-10 mice/group for mRNA expression level measurement, and 3-4 mice/group for flowcytometry. Statistical significance is indicated by *$p$ < 0.05 according to unpaired *t-test* or Mann Whitney test.

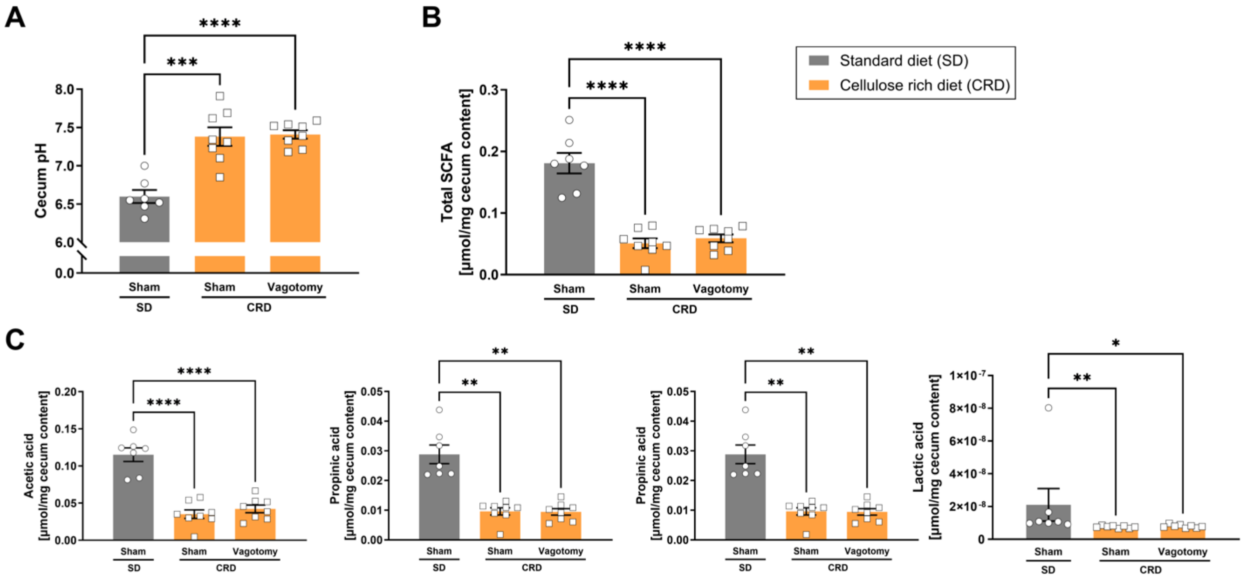


**Supplementary Figure 3: Vagotomy had no effect on the cecum pH and intestinal SCFA concentration.**

Increase of cecum pH (A) and significant decrease of SCFA and lactic acid in cecum content (B, C) induced by chronic CRD consumption has not been altered by vagotomy. Data are expressed as means±SEM of 7-8 mice/group. Statistical significance is indicated by ***$p$ < 0.005, ****$p$ < 0.001 according to unpaired *t-test* or Mann Whitney test.

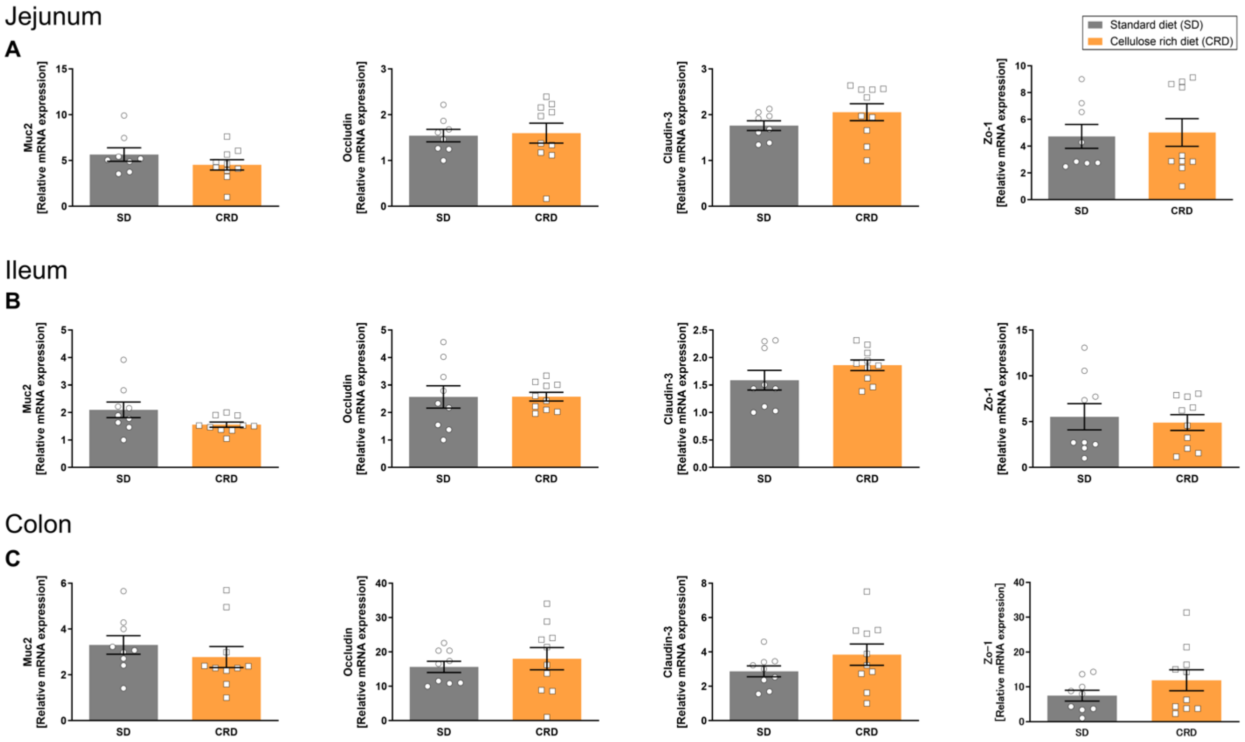


**Supplementary Figure 4: CRD consumption had no effect to the mRNA expression of tight junction genes at any intestinal tissue.**

The mRNA expression levels of Muc2, Occuldin, Claudin-3 and Zo-1 has been evaluated to see the effect of CRD consumption to the intestinal tight junction. Jejunum (A), ileum (B) and colon (C) taken from mice fed each diet for 16 weeks has been used. CRD had no significant effect to the tight junction-related genes, despite of increased permeability (see Figure 4). Data are expressed as means±SEM of 8-9 mice/group.

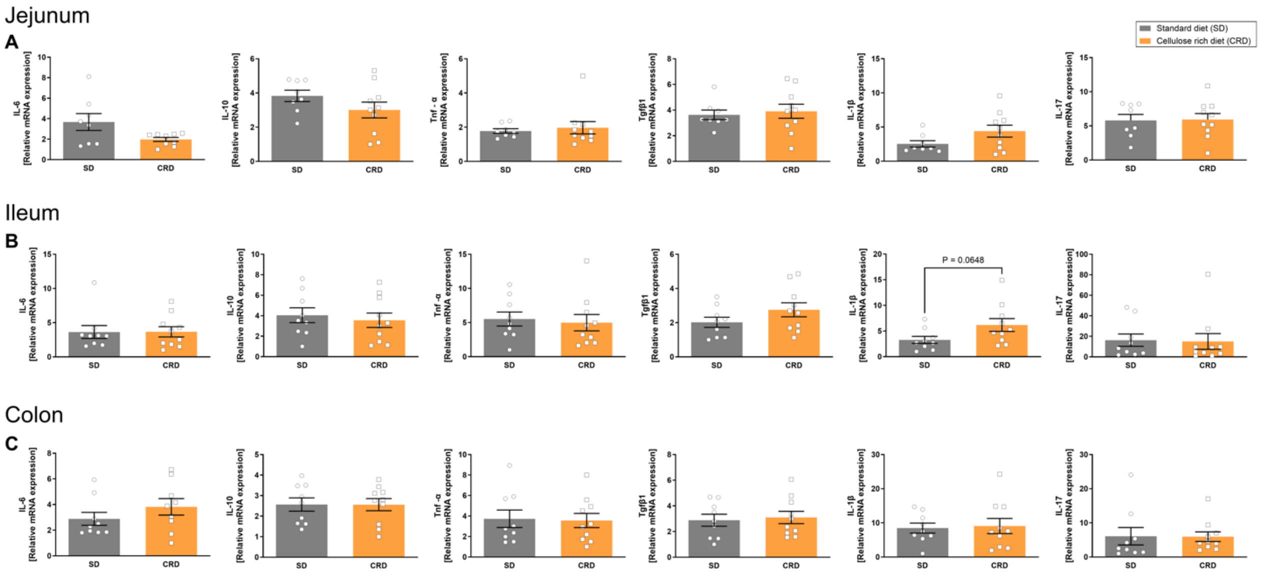


**Supplementary Figure 5: CRD consumption did not affect the mRNA levels of cytokines.**

The mRNA expression levels of pro- and anti-inflammatory cytokines has been evaluated to see the effect of CRD consumption to the intestinal inflammation. Jejunum (A), ileum (B) and colon (C) taken from mice fed each diet for 16 weeks has been used. CRD had no significant effect to any cytokine genes, except the slight increasing tendency on IL-1β level in ileum. Data are expressed as means±SEM of 8-9 mice/group. Statistical significance is indicated according to unpaired *t-test.*

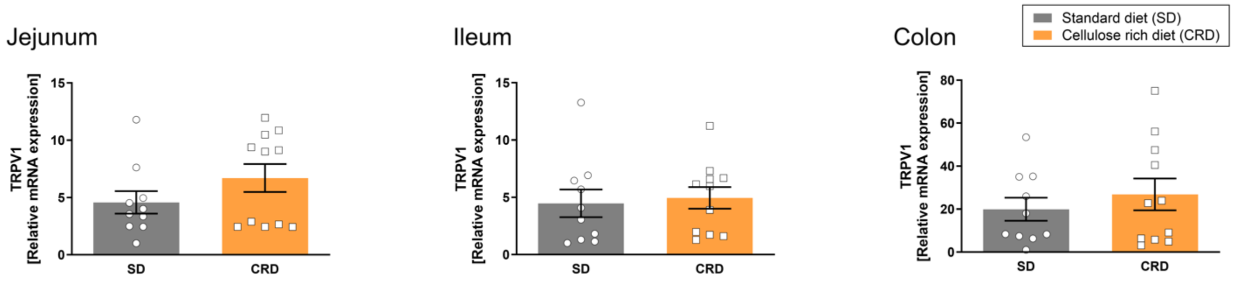


**Supplementary Figure 6: CRD consumption did not modify mRNA expression TRPV1.**

The mRNA expression levels of TRPV1 in intestinal tissue was measured by real-time RT-PCR. Tissues were taken from mice fed either SD or CRD for 8-weeks. Data are expressed as means±SEM of 8-9 mice/group. Statistical significance is indicated by *$p < 0.05$ according to unpaired *t-test*.

**Supplementary Table 1: Primer sequences for RT-PCR.**

| | Forward (5'→3') | Reverse (5'→3') |
|---|---|---|
| D1R | GTAGCCATTATGATCGTCAC | GATCACAGACAGTGTCTTCAG |
| D2R | GCAGCCGAGCTTTCAGGGCC | GGGATGTTGCAGTCACAGTG |
| DAT | GGCTTACAGGACCTCAAAG | TGAACCTCCACTGGTGTCT |
| IL-6 | TGGCTAAGGACCAAGACCATCCAA | AACGCACTAGGTTTGCCGAGTAGA |
| IL-17 | CTCCAGAAGGCCCTCAGACTAC | AGCTTTCCCTCCGCATTGACACAG |
| Tgfβ1 | AGTGTGGAGCAACATGTGGA | GTACAACTCCAGTGACGTCA |
| IL-10 | AGAGAAGCATGGCCCAGAAA | ACACCTTGGTCTTGGAGCTT |
| Tnf-α | GAGCACAGAAAGCATGATCC | CCACAAGCAGGAATGAGAAG |
| IL-1b | TCCATGAGCTTTGTACAAGG | GGTGCTGATGTACCAGTTGG |
| Muc2 | ACCTGGAAGGCCCAATCAAG | CAGCGTAGTTGGCACTCTCA |
| Occludin | ATGTCCGGCCGATGCTCTC | TTTGGCTGCTCTTGGGTCTGTAT |
| Claudin-3 | CAGGGGCAGTCTCTGTGCGAG | GCCGCTGGACCTGGGAATCAAC |
| Zo-1 | TCATCCCAAATAAGAACAGAGC | GAAGAACAACCCTTTCATAAGC |
| TRPV1 | AGCTGCAGCGAGCCATCACCA | ATCCTTGCCGTCCGGCGTGA |
| TRPA1 | TCCGGTCGATCTCAGCAATG | GGCAATGTGGAGCAATAGCG |
| SGLT1 | CCATGGACAGTAGCACCTTGAGC CCC | CTGCCAGGAAGAAGCCTCCAACG GTA |
| 18s rRNA | GGGGAGTATGGTTGCAAAGC | TGTCAATCCTGTCCGTGTCC |

**Supplementary Table 2: RT-PCR settings**

<table>
<tr><td>Reverse transcript</td><td>40℃, 5 minutes</td><td rowspan="2">×1</td></tr>
<tr><td>Pre-PCR</td><td>95℃, 10 seconds</td></tr>
<tr><td>Denaturing</td><td>95℃, 5 seconds</td><td rowspan="2">×40</td></tr>
<tr><td>Primer annealing, extension</td><td>60℃, 34 second<br>95℃, 15 second<br>60℃, 1 minutes</td></tr>
<tr><td>Dissociation Protocol</td><td>Rises to 95°C（0.3℃/seconds）</td><td>×1</td></tr>
</table>

**Supplementary Table 3: Antibodies used in flow cytometry**

| Fluorophores | Cell markers | Concentration | Manufacturer | Cat. |
|---|---|---|---|---|
| **FITC** | Ly6C | 1:200 | BD Pharmingen | 553104 |
| **PE** | CD21/35 | 1:800 | BD Pharmingen | 552957 |
| **PE-CF594** | CD11b | 1:200 | BD Horizon | 562287 |
| **PE-Cy7** | CD11c | 1:100 | BioLegend | 117318 |
| **APC** | CD161/NK1.1 | 1:800 | BioLegend | 108709 |
| **APC-Cy7** | I-A/I-E | 1:400 | BioLegend | 107628 |
| **BV421** | CD5 | 1:200 | BD Horizon | 562739 |
| **V450** | Ly6G | 1:400 | BD Horizon | 560603 |
| **BV510** | CD19 | 1:200 | BD Horizon | 562956 |
